# Cryo-EM analysis of a viral portal protein *in situ* reveals a switch in the DNA tunnel

**DOI:** 10.1101/713933

**Authors:** Oliver W. Bayfield, Alasdair C. Steven, Alfred A. Antson

## Abstract

The portal protein is a key component of many double-stranded DNA viruses, governing capsid assembly and genome packaging. Twelve subunits of the portal protein form a ring with a central tunnel, through which DNA is translocated into the capsid. It is unknown how the portal protein functions as a gatekeeper, preventing DNA slippage, whilst allowing its passage into the capsid through its central tunnel, and how these processes can be controlled by capsid and motor proteins. A cryo-EM structure of a portal protein, determined *in situ* for immature capsids of thermostable bacteriophage P23-45, suggests how domain adjustments can be coupled with a switching of properties of the DNA tunnel. Of particular note is an inversion of the conformation of portal loops which define the tunnel’s constriction, accompanied by a switching of surface properties from hydrophobic to hydrophilic. These observations indicate how translocation of DNA into the viral capsid can be modulated by changes in the properties and size of the central tunnel and how the changing pattern of protein–capsid interactions across a symmetry-mismatched interface can facilitate these dynamic processes.

## Introduction

Tailed bacteriophages constitute the majority of viruses in the biosphere (1, 2) and are a significant component of the human microbiome (3). During assembly, these viruses translocate their genomic double-stranded DNA through a portal protein that occupies a single vertex of an icosahedral capsid. A similar mechanism is employed by the evolutionarily related herpesviruses (4, 5). Structural information about the portal protein is important not only for deducing the mechanism of capsid assembly (6), but also for understanding molecular events associated with genome translocation into preformed capsids (7–9), and genome ejection during infection (10).

Although structures of isolated portal proteins, outwith the native capsid environment, have been determined to near-atomic resolution by X-ray crystallography and cryo-electron microscopy (7, 11–13), a number of observations concerning these structures have yet to be rationalised in the context of the portal’s many functional roles, including: the variable diameter of the central tunnel and flexibility of tunnel loops (7, 11, 12); the symmetry mismatch between the portal and capsid vertex (12-fold versus 5-fold) (9, 11); and the portal’s role in DNA translocation (14, 15). The influence of the properties of the internal tunnel, and how these can be modulated by external factors to coordinate DNA translocation, remains unclear. Cryo-EM studies on mesophilic herpesviruses characterised the shape of the portal protein tunnel in the mature virion and showed how DNA can be locked inside (5, 16). However, there are no detailed structural data on portal proteins *in situ* for unexpanded capsids, primed for DNA packaging. Moreover, it has proven difficult to derive accurate models for tunnel loops of the portal protein, such as in the case of tailed bacteriophage ϕ29, where flexible nature of tunnel loops prevented their observation in structural data (11, 17).

To gain knowledge about the structure of the dynamic DNA tunnel, we utilised a thermostable bacteriophage, P23-45. Thermophilic viruses must package their genomes under extreme temperature, imposing additional challenges compared to their mesophilic counterparts. This *Thermus thermophilus* bacteriophage is one of the few viruses for which conditions for packaging DNA into capsids *in vitro* have been established, where isolated empty capsids were demonstrated to be competent at packaging DNA (18). Previous cryo-EM reconstructions of procapsids (unexpanded) and mature capsids (expanded), in which icosahedral symmetry was imposed, have revealed the extent of conformational changes that the major capsid proteins undergo upon capsid maturation. During the transition, the capacity of the capsid nearly doubles (18). In this study, the structure of the portal protein *in situ*, and analysis of the reconstruction of the unexpanded procapsid without imposing icosahedral symmetry, reveal substantive conformational differences in the structure of the portal protein (18). The most remarkable difference, induced *in situ*, is an inversion in the conformation of tunnel loops of the portal protein. The tunnel loop inversion “switches” the surface properties at the tunnel’s constriction from hydrophobic to hydrophilic and creates a wider opening. These observations indicate the capsid shell plays a role in defining the conformation and properties of the portal protein, modulating DNA translocation into capsid.

## Results

### Structure of the *in situ* procapsid portal

P23-45 procapsids were purified from lysates of infected *Thermus thermophilus* cells (Fig. 1A). The procedures used for cryo-EM data collection and computing the icosahedrally-averaged reconstruction were described earlier (18). The *in situ* structure of the portal protein within the procapsid was determined by localised reconstruction of portal-containing vertices to a resolution of 3.7 Å by averaging around the 12-fold symmetric axis (Table S1, Fig. S1) (19). The portal protein oligomer is a ring of 12 subunits (Fig. 1B, C, Movie S1). Most amino acid side-chains were clearly resolved in the map, enabling construction of a reliable atomic model (Fig. 1D, PDB 6QJT). Comparison with the crystal structure of the portal protein (PDB 6IBG) reveals several differences: notably, in the positions of the Crown and Wing domains and in the conformation of the tunnel loops (Fig. 2, Movie S2). In the *in situ* structure, the C-terminal Crown domain (residues 377–436) is shifted upwards away from the main body by ∼5 Å (Fig. 2A, B), and twisted by ∼13° around the central axis (Fig. 2C, Movie S2). The Wing domain is pivoted ∼8° downwards, towards the Clip (Fig. 2B, Movie S2).

**Fig. 1.**
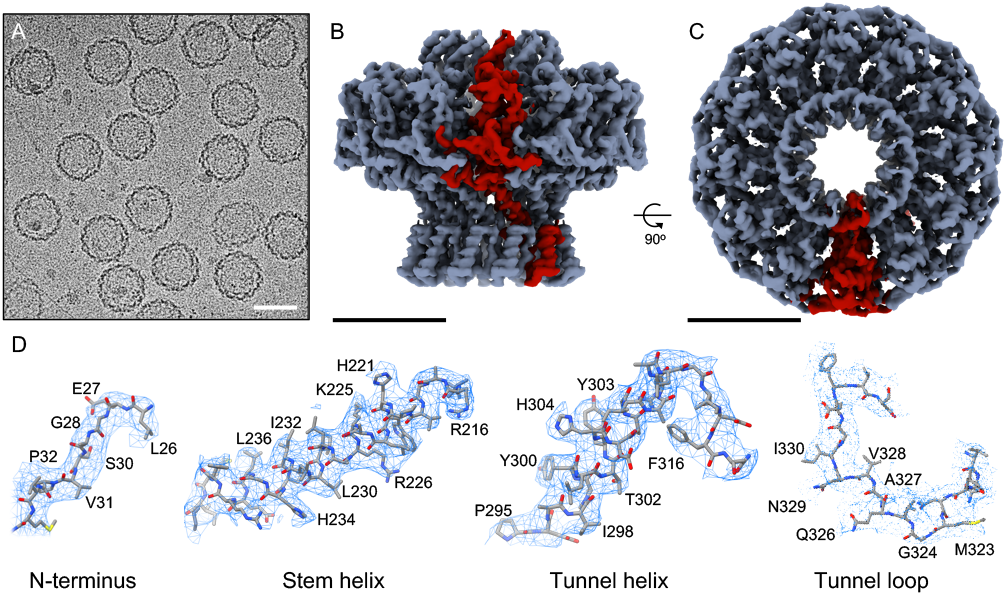
Structure of the portal protein *in situ*. (A) Cryo-electron micrograph of P23-45 procapsids, scale bar 50 nm. (B) Cryo-EM reconstruction map with one subunit coloured red, scale bar 50 Å, and same for (C) but rotated 90°, viewed along 12-fold axis. (D) Regions of the map and corresponding atomic models with residue numbers.

**Fig. 2.**
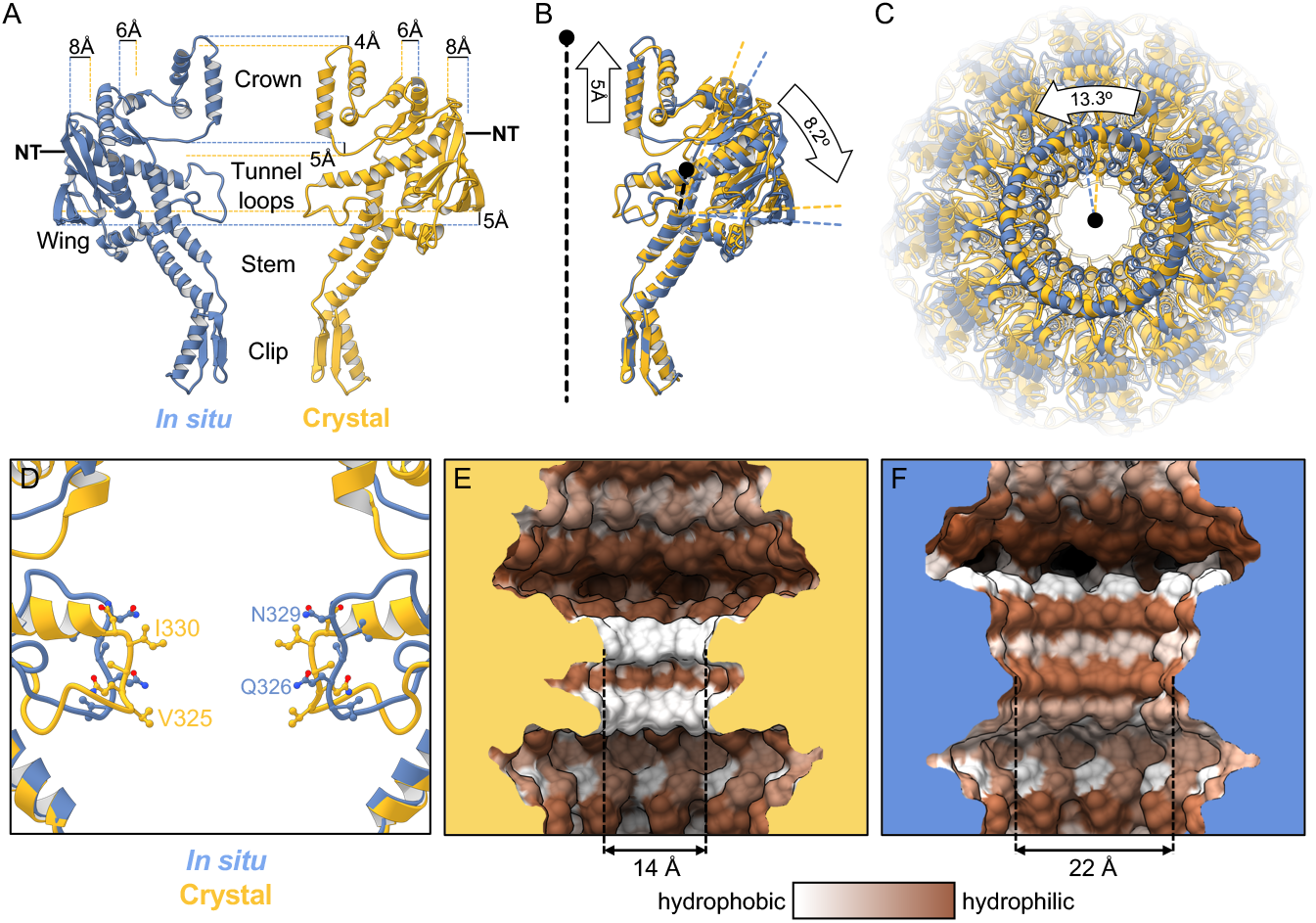
Comparison with the crystal structure. (A) Single subunit of the *in situ* structure is in blue and an apposing chain from the crystal structure is in yellow. (B) Superposition of single subunits, exposing structural differences between the crystal structure and the *in situ* structure. The curved arrow indicates the pivoting of the Wing domain by ∼8° in the *in situ* structure. (C) The two dodecamers over layed, viewed from Crown (top domain in A), along the tunnel axis. Dodecamers are superposed based on residues 26–376 (Clip, Stem, and Wing), revealing a ∼13° rotation of the Crown domain about the tunnel axis. (D) Overlay of *in situ* (blue) and crystal structure (yellow), ribbon diagram, with side-chains shown. (E) Van der Waals surface of crystal structure (PDB 6IBG) showing tunnel loop constricted region, with tunnel colouring by the hydrophobicity KyteDoolittle scale where white is hydrophobic and brown is hydrophilic, and same for (F) but for *in situ* structure (PDB 6QJT). Diameters of most constricted part of tunnels measured from Van der Waals surfaces are shown.

### Differences between the portal conformations in the *in situ* and crystal structures

The most pronounced conformational differences seen in the *in situ* structure are in the tunnel loops (Fig. 2D). The tunnel diameter at its most constricted part is widened by ∼8 Å (Fig. 2E, F). Hydrophobic residues V325 and I330 are no longer exposed to the tunnel as they are in the crystal structure, and are replaced by polar residues Q326 and N329 due to inversion in the tunnel loop conformation (Fig. 2D). Residues 330–335, which protrude into the tunnel and are part of the longest helix in the crystal structure, instead adopt an extended loop conformation *in situ* (Fig. 2D), facilitating the tunnel loop remodelling. These modifications alter the shape and surface properties of the tunnel, which widens and changes from hydrophobic to hydrophilic (compare Fig. 2E and F).

The first N-terminal residue that could be reliably modelled in the *in situ* reconstruction was Leu26 (Fig. 1D), in common with the crystal structure (PDB 6IBG). Mass spectrometry detected N-terminal residues of the portal protein subunits (Fig. S2), indicating that the 25-amino acid N-terminal segment is present in at least some chains, but adopts variable conformations. Although the first residue with a defined conformation points toward the interior of the capsid in P23-45, it cannot be ruled out that the flexible N-terminal segment folds back and contributes to portal-capsid interactions.

### Portal–Capsid interactions

The portal–capsid interface is spacious, with only a small portion of the portal’s outer surface engaged in interactions, predominantly at the height of the portal Wing and Clip (Fig. 3). Residues of the portal Wing involved in these interactions are 185–189 (*β*-hairpin loop) (Fig. 3B). These loops could easily pivot downwards to make closer contact with the capsid inner wall, in select chains (Fig. 3B, C). Such local adjustments would not be resolved in a symmetrically averaged structure, but bridging regions between the capsid and portal in the asymmetric capsid reconstruction suggests local deviations from C12 symmetry are possible. Interactions of the portal Clip likely involve residues K263, E264, K267, N268, Q271, K272, R274, H275 (*α*-helix and adjacent loop within Clip domain), with all points of portal–capsid connection seemingly involving Lysine side-chains of the portal (Fig. 3D). The portal Wing is in close proximity to the T-loop (R337, N338, K340) and N-terminal segment (residues ∼24–30) of the surrounding capsid monomers. The portal Clip domain interacts with the capsid protein P-domain (residues ∼119–127, D357, D358). In common with ϕ29 (11), portal-capsid interactions are mediated by residues of both polar and hydrophobic character. The portal–capsid symmetry mismatch means that only select portal chains make contact with the capsid: these are chains A-C-(E/F)-H-J at the Wing (Fig. 3E), and chains C-E-(G/H)-J-L at the Clip (Fig. 3F).

**Fig. 3.**
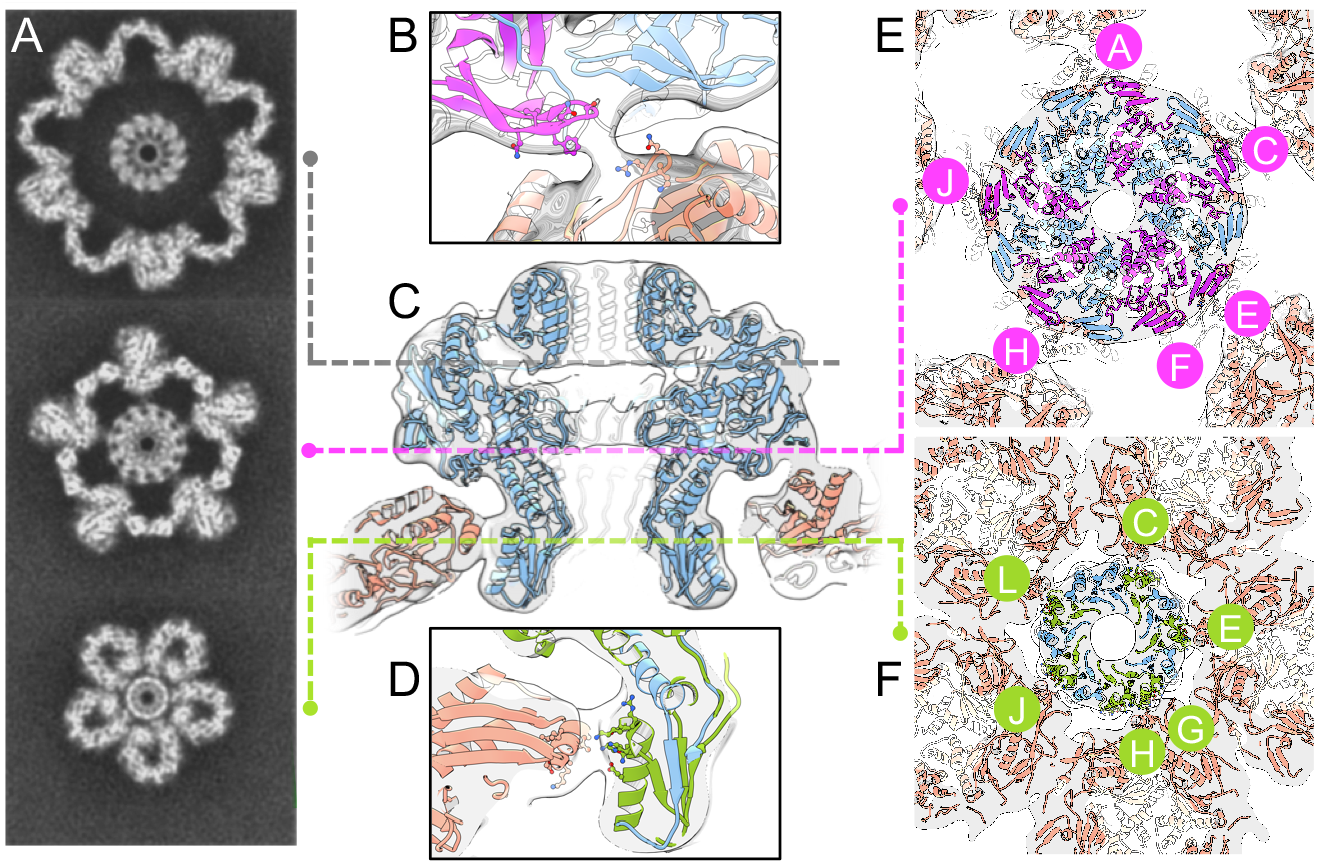
Portal–capsid interactions. (A) Sections through the capsid reconstruction perpendicular to the portal tunnel axis, at heights corresponding to the adjacent panels denoted by dotted lines. (B) Ribbon diagram with sidechains shown for a portal Wing and capsid interacting region. (C) Ribbon diagram of the *in situ* portal protein fitted into the procapsid map. (D) Ribbon diagram with sidechains shown for a portal Clip and capsid interacting region. (E) Interacting regions of the portal Wing with chains labelled clockwise, and (F) of the portal Clip, viewed from the center of capsid (i.e. from the top in panel C). Portal chains making contact with the capsid are coloured in magenta (Wing) and green (Clip).

## Discussion

### Procapsid assembly primes the portal for packaging

The *in situ* structure of the portal protein differs from the crystal structure globally, in alteration to domain positions, and locally, in conformational changes such as the inversion of the tunnel loop. Structural data indicate how the changes on these two levels are linked:

1. Assembly of capsid proteins around the portal stabilises a ∼8° rotational adjustment in Wing domains.
2. As the Wing domain pivots, the C-terminal helix of the Crown domain slides along the Wing surface, facilitating a ∼5 Å shift of the Crown domain towards the capsid centre.
3. This movement of the Crown creates space between the Wing and Crown, which allows remodelling of tunnel loops, facilitated by an unfolding of a 5-residue segment of the long helix within the Wing domain.
4. The loop remodelling “switches” the properties of the tunnel surface from hydrophobic to hydrophilic, causing the tunnel to “open” at its most constricted part.
5. Reversal of the Crown and tunnel movements has the ability to “close” the tunnel.

It is reasonable to assume that the two conformational states, observed in structural studies, reflect energetically preferred states of the portal protein that are utilised during DNA translocation. The switch between the open and closed states, resulting in alteration of surface properties of the internal tunnel may therefore serve a role in the packaging mechanism. The observed conformational differences between the two portal protein states are consistent with the normal mode analysis (18), suggesting a dynamic equilibrium between these two states exists. Analogous conformational changes in central tunnel, involving hydrophobic–hydrophilic “switch” in surface properties, have been proposed to play a key mechanistic role in other systems, for example the GroEL molecular chaperone, where ATP-induced changes facilitate protein refolding (20–22).

It is important to consider how the two portal states are related, and how they may participate in the DNA translocation mechanism. Whereas the ∼22 Å wide, hydrophilic tunnel observed *in situ* would allow the passage of B-form and potentially even the wider A-form DNA (14, 15) into the capsid, the more restrictive tunnel diameter of ∼14 Å observed in the crystal structure require the tunnel loops to protrude towards DNA grooves, involving changes in the tunnel loop conformations (12).

### A model for DNA translocation

Based on structural observations, we propose the following model for DNA translocation (Fig. 4). In this scenario, the tunnel loops engage with DNA to prevent its slipping: modulating the length of packaging dwell periods as the capsid fills, as observed in single-molecule experiments in the ϕ29 system (23–25); and acting to arrest slipping, as observed in single molecule experiments in the T4 system (26). At the start of a packaging cycle, the tunnel is open for DNA translocation, with the tunnel loops widely separated and hydrophilic. During one cycle, it is has been observed that 10 basepairs of dsDNA are translocated for every 4 molecules of ATP hydrolysed (25). After one ATP hydrolysis cycle, when the motor resets and binds new ATP (24, 25), the ATPase must undergo conformational changes during which it is likely to disengage with DNA (27). This poses the risk that the packaged genome could slip—partially or fully—from the capsid; a risk which presumably increases as the pressure inside the head builds. In this instance, the portal tunnel loops can act to negate this risk, constricting to prevent the tendency of DNA to slip out of the capsid. In this model, the portal tunnel is able to switch to a constricted “closed” state when slippage starts to occur. Constriction of the tunnel could be induced by slight slippage of DNA, which would cause downward movement of the Crown domain due to Crown’s coulombic attraction to DNA (DNA “catching” on the Crown). Downward movement of the Crown can have an effect on the tunnel loop conformation, as detailed in points 1 to 5 above. The constricted state could also be induced by an overall downward pressure on the Crown domain, which builds as capsid fills with DNA. The constricted portal tunnel state would be induced more readily as the capsid fills and the internal pressure builds, which could be the cause of dwell times lengthening (23). When more severe slippage occurs, as has been observed in single molecule experiments for bacteriophage T4 (26), this mechanism would arrest slippage, so that normal packaging cycles can resume. The putative role of the tunnel loops in engaging with the DNA is supported by the observations that tunnel loop deletions cause DNA to escape from the capsid in both ϕ29 and T4 phages (28, 29), prior to the tail binding the portal. Overall, this describes a packaging mechanism that is naturally safeguarded against genome loss. At full packaging, DNA can be held in place by constricted tunnel loops and reinforced by tail factors (5, 16).

**Fig. 4.**
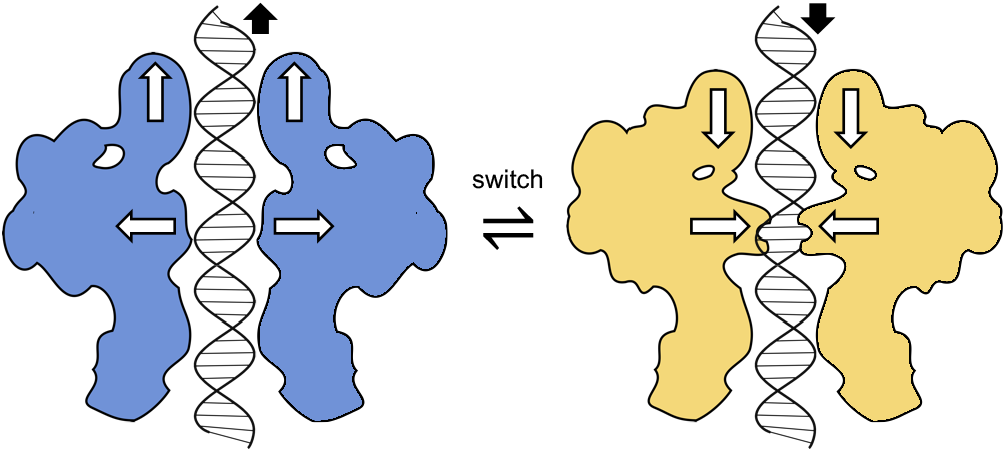
Tunnel constriction by loops. Left, tunnel loops are widely spaced and the internal surface hydrophilic (tunnel “open”); the large terminase is engaged with DNA and converts ATP hydrolysis into mechanical translocation of DNA into the capsid. Right, tunnel loops are constricting the tunnel and extending towards DNA (tunnel “closed”); the large terminase can disengage.

### The portal’s high order of symmetry reconciles a symmetry mismatch

The C12-symmetric portal is accommodated in a C5-symmetric penton cavity at one capsid vertex, despite the attendant symmetry mismatch. The ∼8° rotational adjustment of the Wing position, bringing it closer to the capsid wall, may assist in forming close portal–capsid contacts, whereas the portal’s Clip external diameter is already well matched to the aperture of capsid’s penton tunnel so that close interactions can be made. However, the symmetry mismatch creates the problem in that the same pairs of interacting residues at the portal–capsid interface are not consistently aligned around all subunits, and could be offset by as much as ∼2 nm in the case of P23-45. This misalignment occurs both at the portal Wing and at the Clip, where the portal and capsid make contact in the asymmetric capsid reconstruction. The sparsity of connected portal–capsid regions indicates that the total surface area of interaction is small, and the residues involved in such interactions are hence also restricted in number and positioning.

The high order of symmetry of the portal helps to mitigate these problems. Its 12-fold symmetry is advantageous in that regions of the portal which can interact with the capsid are repeated at a correspondingly high frequency, which reduces the distance between the mismatched interacting residues. The remaining distance can easily be closed by pivoting of flexible loops towards the capsid, such as at the *β*-hairpin loops of the portal Wing (residues 185–189). These loops are in equivalent positions in ϕ29 (17). As a result, only minimal, localised deviations from ideal C12 symmetry are needed to make interactions with the capsid. The portal can therefore utilise the same few residues to interact around its circumference, which contrasts with the situation that would exist if the portal possessed C6, C3, or other lower symmetries matching that of tail components. The symmetry mismatch of the interaction is a general feature amongst all tailed bacteriophages and related viruses, including herpesviruses (5, 16). In the case of bacteriophage ϕ29 prohead (8, 17), one of the structural roles of the pRNA appears to be equivalent to that of the capsid protein P-domain, in interacting with the outside of the portal Clip, with the ϕ29 capsid protein Pdomain instead making contact with an N-terminal segment of the portal protein.

At the interface between the portal and capsid vertex, with respective C12 and C5 symmetries, interactions will repeat with a periodicity of 360°/60 = 6°, as similarly suggested by Hendrix prior to the determination of portal structures (30). Rotation of the portal with respect to the capsid by only 6° would therefore create an equivalent global register, with rotations less than 6° generating non-superposable registers of the whole capsid particle (Movie S3). Different portal–capsid registers will have different energies of interaction, and hence equivalent angular registers are expected to be energetically equivalent. A comparable symmetry mismatch is observed between the portal and internal core of bacteriophage T7 (C12 *versus* C8) (31), where the mismatched interactions may facilitate the detachment of core proteins. As neither detachment of the portal, nor its rotation with respect to the capsid (32), seem likely to play a role in capsid maturation, the effect of symmetry mismatch in the capsid vertex is to permit flexibility at the portal–capsid interface, allowing the portal and capsid to undergo independent conformational changes, whilst ensuring stable interaction of the portal protein with the tail.

## Conclusions

Accommodation of the portal protein dodecamer in the procapsid involves conformational adjustments. Interaction between the portal and the capsid alters relative positions of domains, in particular the Wing and Crown, and causes remodeling in the tunnel loops that define the most constricted part of the axial tunnel. The unique conformation of the portal *in situ* demonstrates that the capsid plays a role determining portal conformation, allowing DNA to pass through the tunnel, whilst the portal has the ability to modulate packaging activity and slippage by switching its tunnel properties so that it can engage and disengage with DNA. Whilst portal proteins across all double-stranded DNA viruses may deviate from the classical domain arrangement observed in P23-45, all such viruses face the same basic challenge of safeguarding against genome loss. With regards to portal-capsid interactions, the adoption of 12-fold symmetry by the portal, rather than a symmetry matching that of the capsid vertex, is likely a consequence of the independent evolution of head and tail assemblies, which has selected the matching of symmetries between the portal (12-fold) and the tail (6-fold). This study posits that the problem of mismatched portal–capsid interactions is resolved by the high number of subunits constituting the portal protein, which minimises distances between interacting regions across a spacious interface.

## Methods

### Cryo-EM data processing and model building

From 38,044 extracted particles used in the reconstruction of the unexpanded icosahedral capsid (EMD-4447) (18), subparticles centred on each vertex were extracted from each capsid particle, and were aligned on the z-axis (19). After 3D classifications without imposing symmetry or changing orientations in RELION (33), a class containing 10,025 particles exhibiting clear portal features was selected for subsequent 3D refinement in RELION, with C12 symmetrical averaging. The atomic model was built using the crystal structure PDB 6IBG as a starting model, with modification to domain positions and to individual amino acids, including side chain conformations, introduced in Coot (34). Cycles of model rebuilding were followed by real-space refinement in Phenix (35). Resolution was assessed using the gold-standard method using the FSC 0.143 criterion. Refinement and model statistics are presented in Table S1. Rendering of figures and structure analyses were performed in UCSF Chimera (36).

### Liquid Chromatography–Mass Spectrometry

Capsids of P23-45 in the unexpanded state were purified as previously described (18), and digested with enzyme Glu-C, followed by liquid chromatography tandem mass spectrometry. A 20 µl aliquot (20 µg protein) was reduced with DDT and alkylated with iodoacetamide. Digestion was performed for ∼18 hours at 37 °C using sequencing grade Glu-C (Promega). Peptides were analysed by nanoHPLC-MS/MS over a 65 min acquisition with elution from a 50 cm C18 PepMap column onto an Orbitrap Fusion Tribrid mass spectrometer via an Easy-Nano ionisation source. LC-MS/MS chromatograms were analysed using PEAKS-Studio X (37). Peaks were picked and searched against the combined *Thermus thermophilus* and Thermus phage P23-45 proteomes. Database searching required Glu-C specificity with one site of non-specificity per peptide identity allowed. Expected cleavage is C-terminal to Glu, a lower rate of cleavage C-terminal to Asp is also known to occur. Peptide matches were filtered to achieve a false discovery rate of <1%.

### Data deposition

Deposition of the cryo-EM reconstruction (EMD-4567) and atomic coordinates (PDB 6QJT) were made using the wwPDB (www.wwpdb.org).

## ACKNOWLEDGEMENTS

The authors thank the Astbury Centre, University of Leeds, for assistance with cryo-EM data collection and helpful discussion; Adam Dowle at the Biology Technology Facility, University of York, for assistance with mass spectrometry. This work was supported by Wellcome Trust–National Institutes of Health Studentship 103460 (to O.W.B.), Wellcome Trust Grant 206377 (to A.A.A.), and by the Intramural Research Program of National Institute of Arthritis and Musculoskeletal and Skin Diseases (A.C.S.).

