## Supplementary for "Cryo-EM analysis of a viral portal protein *in situ* reveals a switch in the DNA tunnel"

#### Figures

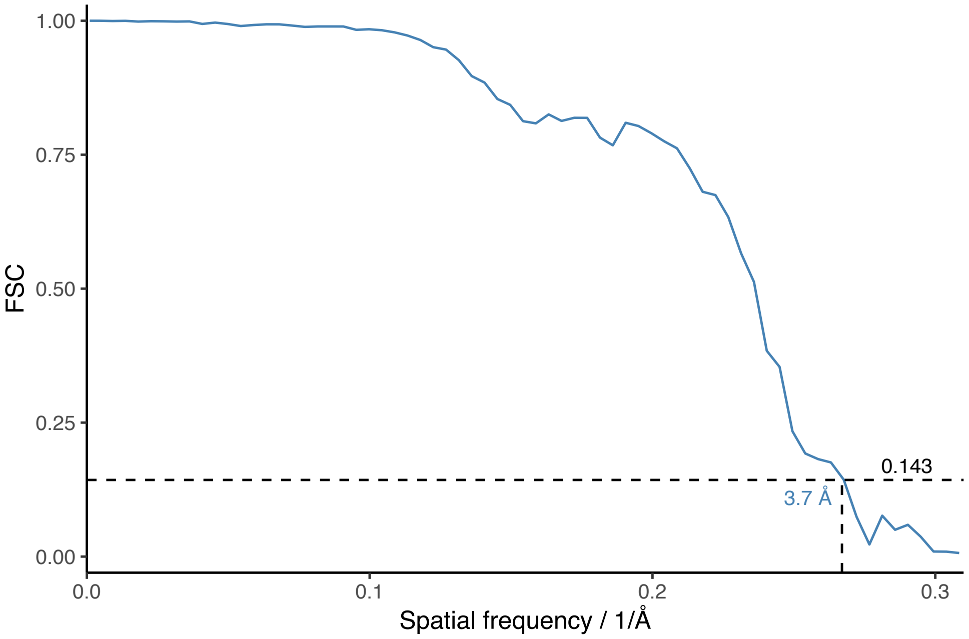


**Fig. S1.** FSC curve for the portal protein reconstruction. Fourier Shell Correlation is plotted as function of spatial frequency. Dotted lines denote resolution estimate at FSC=0.143 according to the gold-standard method.

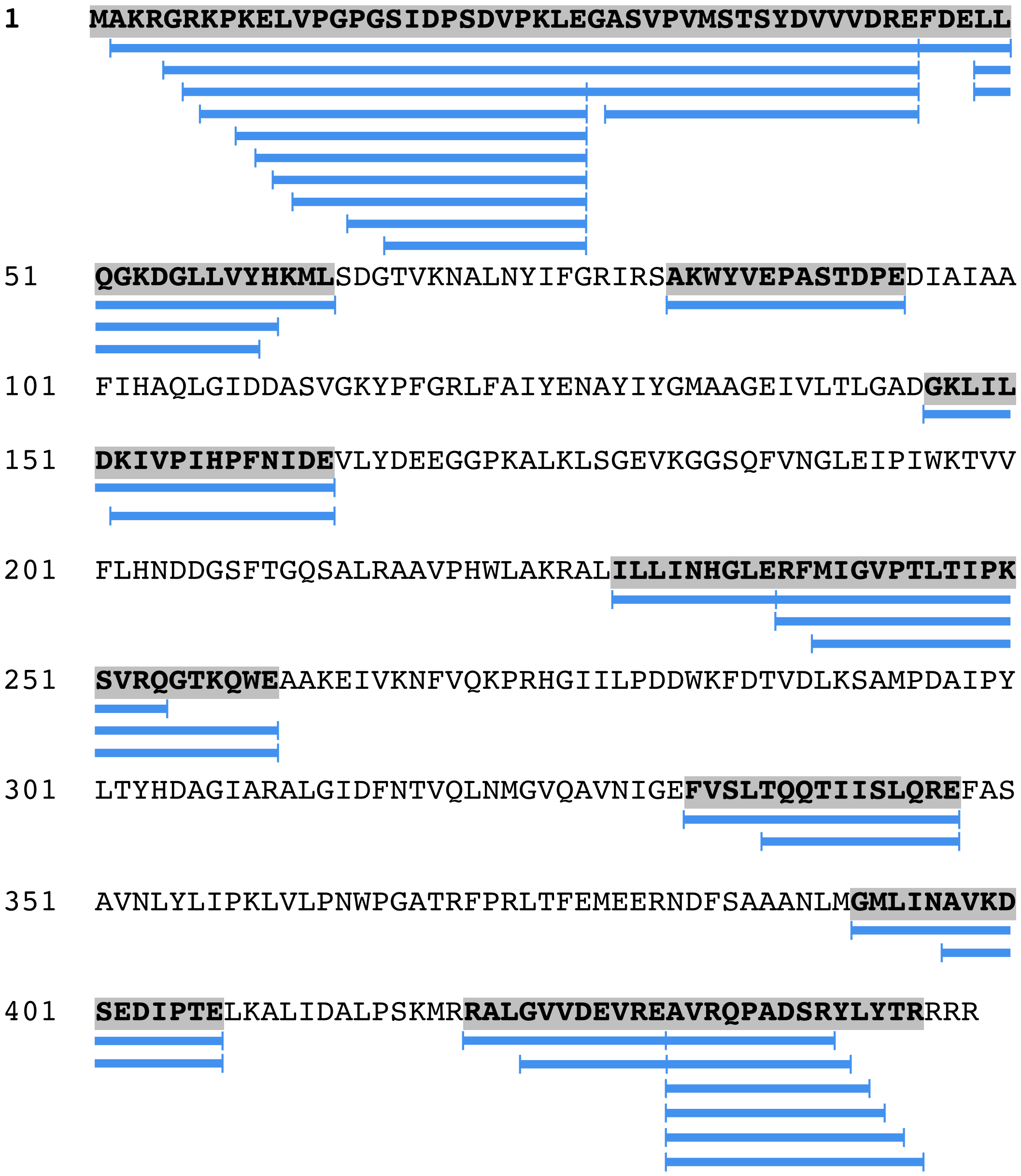


**Fig. S2.** Mass spectrometry analysis of the portal protein from unexpanded capsids. Blue bars beneath the sequence denote regions for which peptides were detected.

### Tables

**Table S1.** Cryo-EM Data Collection and Refinement Statistics.

| | |
|---|---|
| **Data collection** | |
| Microscope | Titan Krios |
| High Tension / kV | 300 |
| Pixel size, unbinned / Å | 1.065 |
| Spherical aberration / mm | 2.7 |
| Nominal magnification | 75000 |
| Nominal defocus / μm | 0.5–2.5 |
| Detector (mode) | Falcon 3EC (integrating) |
| Accumulated dose / eÅ$^{-2}$ | 99 |
| **Refinement and model statistics** | |
| Symmetry | C12 |
| Resolution (FSC 0.143) | 3.74 |
| Map-to-model correlation | 0.835 |
| MolProbity score | 1.36 |
| EMRinger score | 2.56 |
| RMS deviations | |
| Bond lengths / Å | 0.005 |
| Bond angles / ° | 0.842 |
| Ramachandran plot / % | |
| Favored | 94.62 |
| Allowed | 5.38 |
| Outlier | 0.00 |

#### Movies

**Movie S1.** Reconstruction of the *in situ* portal. Surface rendering, first viewed perpendicular to the tunnel axis, then viewed along the tunnel axis.

**Movie S2.** Morph between the *in situ* structure (first) and crystal structure (second). Ribbon diagram, first viewed perpendicular to the tunnel axis, then viewed along the tunnel axis, then rotated back to initial view with two apposing chains displayed.

**Movie S3.** Portal–capsid registers. One-degree step change in portal register (inner 12-fold circle) with respect to capsid vertex (outer 5-fold circle), beginning with “$0^o$”. Portal register “$6^o$” is superposable on register “$0^o$” by $144^o$ rotation of the whole capsid (i.e. rotating both inner and outer circles together).
